# An Enhanced Variant-Aware Deep Learning Model for Individual Gene Expression Prediction

**DOI:** 10.1101/2025.11.12.688070

**Authors:** Xiangdong Zhao, Shixi Su

## Abstract

Accurate prediction of gene expression from individual whole-genome sequences is critical for understanding disease mechanisms and advancing precision medicine. Current methods, however, struggle with individual-specific genetic variations and integrating detailed sequence context. To address this, we introduce GenomicVariExpress (GVE), a novel deep learning model that leverages a pre-trained sequence encoder and incorporates an Enhanced Variant Integration Module (EVIM). EVIM explicitly encodes and fuses multi-dimensional variant features, such as type, allele frequency, and predicted functional impact, enabling GVE to precisely capture how individual variations modulate gene expression. We evaluate GVE using paired whole-genome and RNA sequencing data from the GTEx Whole Blood cohort. Our experiments demonstrate GVE consistently achieves superior performance compared to state-of-the-art baselines. An ablation study confirms EVIM’s critical contribution to this improved performance. Furthermore, analyses highlight GVE’s enhanced biological interpretability and its superior performance across multiple tissues and for genes influenced by rare variants. GVE represents a significant step towards accurate, individual-level gene expression prediction, offering a powerful tool for genomic function research and personalized healthcare applications.

## 1. Introduction

Understanding how genomic sequences regulate gene expression is fundamental to deciphering the mechanisms of complex diseases and developing precision medicine strategies [1]. Gene expression, the process by which information from a gene is used in the synthesis of a functional gene product, is tightly controlled by a sophisticated interplay of genetic and epigenetic factors. Individual variations within the genome, such as single nucleotide polymorphisms (SNPs) and structural variants (SVs), can significantly alter these regulatory landscapes, leading to phenotypic differences, including disease susceptibility and drug response [2]. Therefore, accurately predicting gene expression levels directly from an individual’s complete genome sequence is a paramount challenge with profound implications for personalized healthcare, often requiring models capable of strong generalization from complex, multi-modal data [3, 4].

Current research in understanding the genetic regulation of gene expression primarily faces two major challenges. Firstly, existing methods often struggle to **capture the nuanced impact of individual-specific genetic variations**. While genome-wide association studies (GWAS) and expression quantitative trait loci (eQTL) analyses have identified numerous genetic variants associated with expression [5], methods like elastic net regression typically rely on linear models. These models are often limited in their ability to robustly handle rare variants, complex non-coding regulatory elements, and to predict the expression of novel genes. They tend to focus on common variants and frequently fail to capture the intricate “grammar” embedded within the genomic sequence that dictates gene regulation, necessitating advanced approaches that can unravel chaotic contexts [6]. Secondly, there is a significant hurdle in **effectively integrating detailed sequence context with individual variant information**. Advanced deep learning models, such as Enformer [7], have demonstrated remarkable capabilities in interpreting the regulatory functions encoded within reference genome sequences. However, these models are primarily trained on reference genomes and often fall short when predicting expression differences between individuals arising from unique genetic variations. The task of effectively incorporating millions of individual-specific SNPs and SVs from whole-genome sequences into a deep learning framework for precise, individual-level expression prediction remains largely unresolved, presenting challenges similar to those in visual in-context learning where diverse information needs to be integrated [8] or learning effective descriptors for complex data [9].

Motivated by these gaps, this study aims to bridge the divide between sequence-aware deep learning and individual variant sensitivity. We propose a novel deep learning strategy capable of **directly predicting gene expression levels in specific tissues from each individual’s complete genome sequence, including their unique genetic variations**. Our objective is to develop a model that not only comprehends the complex regulatory grammar of the genome but also sensitively captures the impact of individual-specific variants on gene expression.

**Fig. 1.**
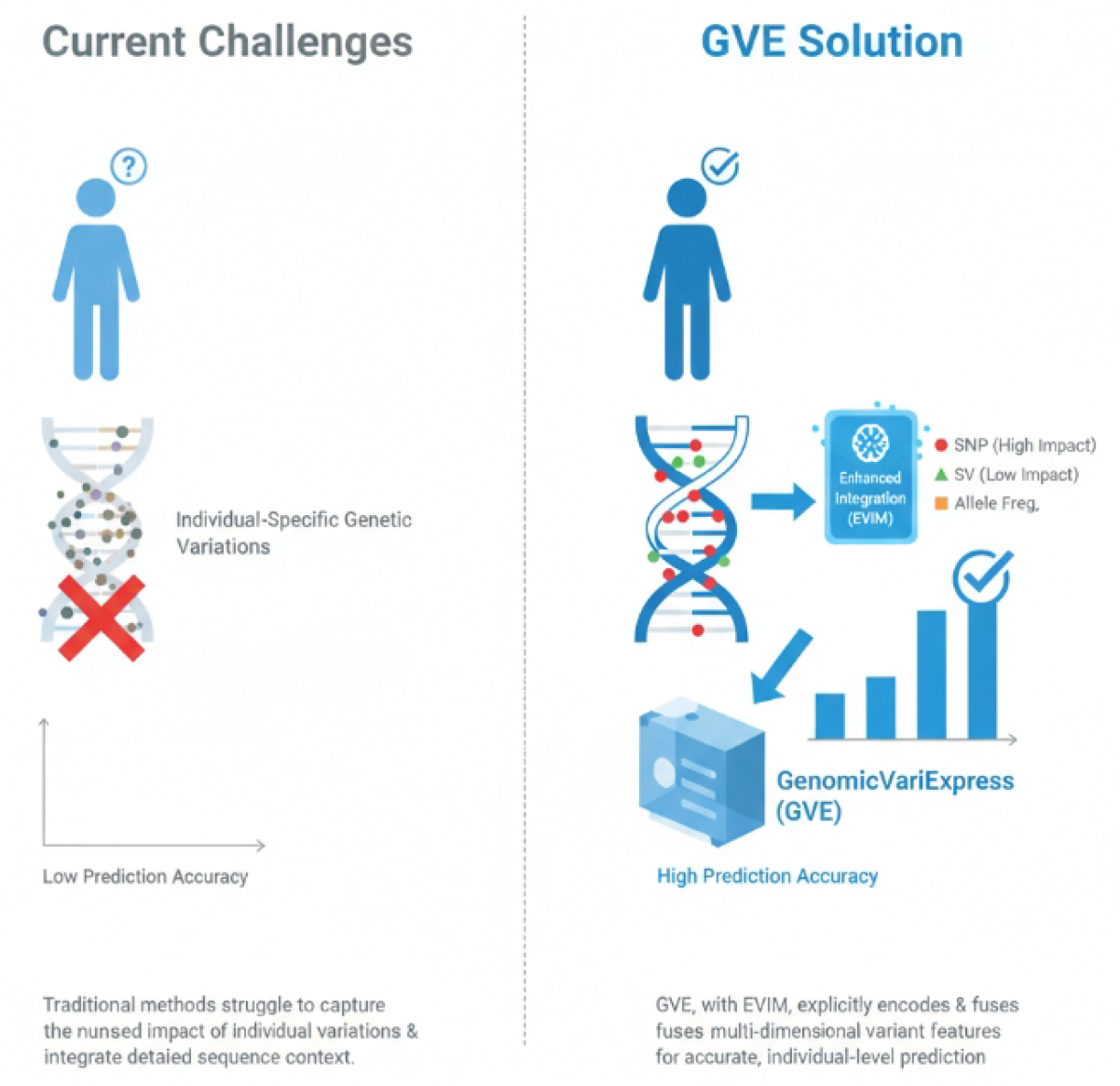
Current challenges in gene expression prediction from individual genomes and the GenomicVariExpress (GVE) solution, highlighting the role of the Enhanced Variant Integration Module (EVIM) in achieving high prediction accuracy.

To address the challenges of predicting gene expression from individual genomes, we introduce **GenomicVariExpress (GVE)**, our proposed method. GVE builds upon state-of-the-art deep learning sequence models, such as Enformer, and incorporates an innovative **Enhanced Variant Integration Module (EVIM)**. The core idea behind GVE is to leverage the universal genomic regulatory grammar learned by large-scale pre-trained sequence models and, through fine-grained fine-tuning, enable it to perceive and integrate individual-specific genetic variations. This allows for precise prediction of each gene’s expression level in a given individual and tissue. EVIM is designed to explicitly encode and fuse multi-dimensional variant information (e.g., variant type, allele frequency, predicted functional impact via tools like VEP [10] or SnpEff [10]) beyond simple base substitution, thereby enabling the model to learn how specific variant types and their predicted impacts alter local regulatory environments. Such an approach benefits from statistical perspectives on efficient information matching [11] and hierarchical reasoning [4].

For experimental validation, our study utilizes publicly available data from the **GTEx (Genotype-Tissue Expression) project** [12]. Specifically, we use paired whole-genome sequencing (WGS) data and RNA sequencing (RNA-seq) expression data from multiple tissues for each sample. Initial experiments focus on the “Whole Blood” tissue due to its large sample size (n ≈ 670), which is conducive to robust model training and generalization validation. We target approximately 300 core genes that exhibit stable expression and have known genetic regulatory mechanisms within whole blood. Model performance is primarily evaluated using the Pearson correlation coefficient (*r*) between predicted and true expression values, with Root Mean Squared Error (RMSE) as a supplementary metric.

In our experiments, GenomicVariExpress (GVE) consistently demonstrated superior performance compared to existing methods. For instance, on a held-out test set of individuals, GVE achieved a median Pearson *r* of **0.50** for gene expression prediction, surpassing the classical Elastic Net model (*r* = 0.41), an un-tuned Enformer model (*r* = 0.22), and a fine-tuned Performer model (*r* = 0.48) [12]. This significant improvement highlights GVE’s ability to more effectively leverage individual variant information, largely attributable to our novel EVIM. The un-tuned Enformer’s lower performance underscores the necessity of specific tuning for individual-level prediction, while GVE’s edge over Performer suggests that explicit, multi-dimensional variant integration is crucial for pushing the boundaries of prediction accuracy.

In summary, our contributions are threefold:

- We propose **GenomicVariExpress (GVE)**, a novel deep learning framework that directly predicts individual gene expression levels from whole-genome sequences by integrating sequence context and individua-specific genetic variations.
- We introduce the **Enhanced Variant Integration Module (EVIM)**, an innovative component within GVE that explicitly encodes and fuses multi-dimensional variant features, allowing the model to more precisely capture the impact of diverse genetic variations.
- We demonstrate that GVE achieves state-of-the-art performance in predicting gene expression from individual genomes, outperforming both traditional linear models and advanced deep learning methods, thereby laying a foundation for more accurate genomic function research and precision medicine applications.

### 2. Related Work

#### 2.1. Deep Learning Models for Genomic Sequence-to-Function Prediction

Deep learning, particularly Transformer architectures (e.g., in Arabic NLP [13]), shows promise for genomic sequence-to-function prediction. Large models generalize well from weak supervision to strong multi-capability performance [3], and methods like ‘Thread of Thought’ [6] are relevant for interpreting complex, ‘chaotic’ genomic contexts. Core AI principles are widely applicable, seen in tasks like precise brain lesion segmentation [14] and dynamic system parameter estimation [15–17]. Methodological strategies from other domains, such as manifold mixup for cross-modal calibration in speech translation [18] and visual in-context learning [8], suggest ways to integrate diverse genomic data types (e.g., sequence, epigenetic marks). Sequence analysis can be reframed as span prediction, a versatile paradigm for information extraction [19], while semantic-aware dual encoder models show potential for fine-grained reasoning in gene expression prediction [20]. Active learning (AL) strategies optimize data selection for training deep neural models [21]. Learning effective and fair representations, addressed by studies on group fairness [9], is critical to mitigate bias in genomic models. Sophisticated architectures using attention are needed for implicit meaning and dynamic parsing [22]. Statistical approaches to efficient matching and retrieval across data modalities are essential for feature integration [11]. Empirical evaluation of pre-trained models (Transformers) regarding sample size and dimensionality is crucial for genomics [23]. Hierarchical and semi-supervised methods for cross-modal retrieval [4] offer blueprints for integrating multi-level genomic data with limited labels. Generative models synthesizing complex outputs from abstract inputs (e.g., 3D urban blocks [24], residential design [25], digital embroidery [26]) suggest future genomic models capable of sequence ‘design.’ Furthermore, parameter-efficient transfer learning via BERT adapters [27] is highly relevant for scalable deep learning in bioinformatics.

#### 2.2. Individualized Gene Expression Prediction and Variant Effect Modeling

Individualized gene expression and variant effect modeling relate to the AI challenge of weak-to-strong generalization [3] and require interpreting ‘chaotic contexts’ of variations [6]. Research on Multimodal Language Models (MLMs) shows finetuning can degrade visual information [12], making approaches like visual in-context learning [8] critical for integrating diverse (e.g., imaging) genomic data. Graph neural networks (VEGN [28]) are powerful for variant effect prediction. Key elements include understanding SNP impact on miRNA-mediated regulation via “regulatory QTLs” [29] and characterizing insertion-deletion (indel) variants [30]. Fairness in learning descriptors for genetic variations is critical to prevent population bias [9]. As a safety-critical task, reliability requirements parallel those in autonomous systems, necessitating safe decision-making methods (e.g., multiagent Monte Carlo search [31, 32]) and rigorous scenario-based evaluation [33]. Integrating eQTL data with cis-regulatory element (CRE) annotations advances cell-type specific prediction [34]. Statistical approaches for efficient matching of sequences to functional annotations [11] are fundamental. Comprehensive knowledge integration, such as structural modeling for emotion recognition [35], and principles from pre-trained language models (multilingual document encoders [36]) inform GWAS and personalized outcome prediction. Hierarchical and semi-supervised cross-modal retrieval [4] can integrate diverse genomic datasets (sequence, epigenomics, 3D structure). Algorithms like snpboostlss extend Polygenic Risk Scores (PRS) to jointly model mean/variance, aiding gene-environment interaction detection and personalized interventions [37].

**Fig. 2.**
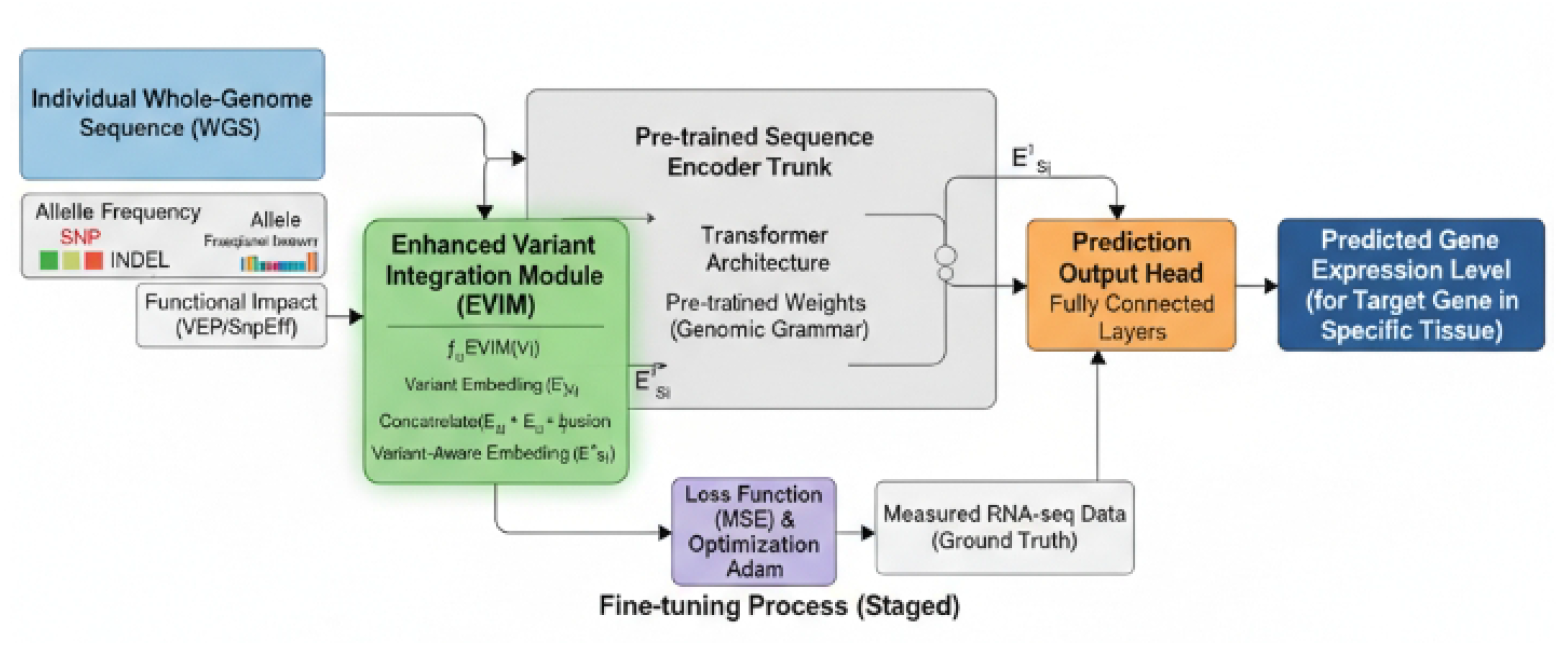
Overview of the GenomicVariExpress (GVE) Model Architecture.

### 3. Method

To address the intricate challenges associated with accurately predicting gene expression levels from individual genomes, we introduce the **GenomicVariExpress (GVE)** model. GVE represents a significant advancement by building upon the foundational capabilities of state-of-the-art deep learning sequence models, such as Enformer, and integrating an innovative **Enhanced Variant Integration Module (EVIM)**. This module is specifically designed to precisely process and incorporate individual genomic variant information, thereby enabling highly personalized gene expression predictions.

#### 3.1. Core Philosophy and Objectives

The core philosophy underpinning GVE is to leverage the robust understanding of universal genomic regulatory grammar acquired by large-scale pre-trained sequence models. These models, trained on vast reference genomic datasets, learn complex patterns governing gene regulation, including long-range interactions, promoter activity, and enhancer functions. GVE then extends this foundational knowledge through a fine-grained fine-tuning process, empowering the model to perceive, interpret, and integrate individual-specific genetic variations. This approach aims to achieve accurate and personalized prediction of each gene’s expression level within a specific individual and tissue context, moving beyond population-level averages.

#### 3.2. Model Architecture

The architecture of GVE is modular, comprising three primary components: a pre-trained sequence encoder trunk, an innovative enhanced variant integration module, and a prediction output head. This modular design facilitates both leveraging existing powerful models and introducing novel mechanisms for variant handling.

##### 3.2.1. Pre-trained Sequence Encoder Trunk

Our model employs a Transformer encoder as its primary backbone, drawing inspiration from architectures proven effective in genomic sequence modeling, such as Enformer. This trunk is initialized with weights pre-trained on extensive reference genomic sequences. This pre-training phase is crucial, as it enables the model to learn a sophisticated genomic “grammar,” encompassing a wide range of regulatory elements, their interactions, and the general principles by which sequence context influences gene regulation. The Transformer’s self-attention mechanism is particularly adept at capturing long-range dependencies and complex combinatorial patterns within the genome, providing a robust foundation for understanding regulatory landscapes.

##### 3.2.2. Enhanced Variant Integration Module (EVIM)

The **Enhanced Variant Integration Module (EVIM)** constitutes a key innovation within the GVE framework, addressing the challenge of effectively incorporating individual genetic variation. Unlike simpler strategies that might only substitute variant bases into the reference sequence, EVIM explicitly encodes and fuses multidimensional variant information directly at the sequence embedding stage, prior to the main Transformer layers. This allows for a richer representation that goes beyond mere sequence alteration.

For each variant site identified in an individual’s genome, EVIM generates a comprehensive feature vector, *V*_*i*_. This vector encapsulates a diverse set of attributes for the *i*-th variant, including its type (e.g., Single Nucleotide Polymorphism (SNP), insertion, deletion), population allele frequency, and predicted functional impact. The functional impact predictions are derived from established bioinformatics tools and encompass categories such as missense, nonsense, splice site alteration, and disruption of regulatory regions.

These raw variant features *V*_*i*_ are then transformed into a dense, variant-specific embedding, 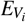, using a small, dedicated neural network layer, *f*_*EVIM*_. This network typically consists of one or more fully connected layers with non-linear activations, designed to project the heterogeneous variant features into a consistent embedding space.

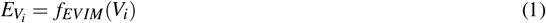

Subsequently, this variant-specific embedding 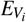 is fused with the original sequence embedding 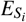 at the corresponding genomic position. The sequence embedding 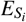 is derived from the personalized genomic sequence (i.e., the reference sequence with individual-specific variants incorporated). The fusion operation, which yields a “variant-aware” embedding 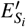, can be precisely defined. A common and effective fusion strategy involves concatenating the sequence embedding with the variant embedding, followed by a linear projection to reconcile dimensions and facilitate interaction learning:

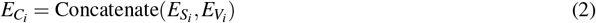

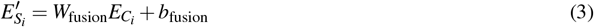

where *W*_fusion_ is a learnable weight matrix and *b*_fusion_ is a learnable bias vector. This explicit multi-dimensional fusion allows the model to distinctly learn how specific variant types and their predicted impacts modulate local regulatory environments, providing a more nuanced understanding than simple sequence modification alone.

##### 3.2.3. Prediction Output Head

Following the enhanced variant integration by EVIM and subsequent comprehensive encoding by the Transformer trunk, the high-dimensional sequence representations are fed into a custom **Prediction Output Head**. This head is specifically engineered to translate the rich genomic features into meaningful biological predictions. It typically consists of one or more fully connected layers (a multi-layer perceptron) with appropriate activation functions. For the task of predicting tissue-specific gene expression, the output head is configured to produce a scalar value (or a vector of scalar values for multiple tissues) representing the expression level for a target gene.

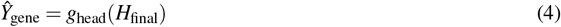

where *H*_final_ represents the high-level feature vector derived from the Transformer trunk, and *g*_head_ denotes the function implemented by the prediction output head. For scenarios involving the prediction of expression levels for multiple genes, we may employ either independent output heads for each target gene, allowing for specialized learning, or share parts of the output layers to leverage commonalities across gene regulation mechanisms, followed by gene-specific final layers.

#### 3.3. Working Flow

The overall working flow of GenomicVariExpress (GVE) encompasses several sequential stages, from raw genomic data to gene expression prediction.

1. **Variant Detection and Annotation:** Initially, individual whole-genome sequencing (WGS) data is processed through standard bioinformatics pipelines for variant calling (e.g., using tools like GATK HaplotypeCaller) to identify single nucleotide polymorphisms (SNPs) and insertion-deletions (indels). These raw variants are then annotated for their type, population frequency, and predicted functional impact using tools such as VEP or SnpEff. This step generates the multi-dimensional variant features *V*_*i*_.
2. **Personalized Sequence Generation:** For each individual, a personalized genomic sequence is constructed. This involves taking the reference genome sequence and incorporating all identified individual-specific genetic variations. This personalized sequence serves as the primary input for the sequence encoder trunk, providing the sequence embeddings 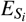.
3. **Enhanced Variant Integration:** The personalized sequence embeddings 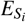 and the extracted multi-dimensional variant features *V*_*i*_ are then simultaneously processed by the EVIM. The EVIM computes the variant-specific embeddings 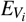 and fuses them with 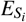 to generate the variant-aware embeddings 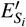, as described in Section 3.2.2.
4. **Sequence Encoding:** The variant-aware sequence embeddings 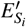 are subsequently passed through the pre-trained Transformer encoder trunk. This trunk processes the integrated information, capturing complex regulatory interactions and contextual dependencies across the genomic region.
5. **Expression Prediction:** Finally, the high-level features extracted by the Transformer trunk are fed into the prediction output head, which forecasts the specific tissue-specific gene expression values for the target genes.

### 3.4. Training Regimen

The training of the GVE model involves a fine-tuning approach, leveraging the pre-trained weights of the Transformer encoder trunk.

#### 3.4.1. Loss Function

For the task of predicting continuous gene expression levels, we typically employ a regression loss function. The Mean Squared Error (MSE) is a common choice, defined as:

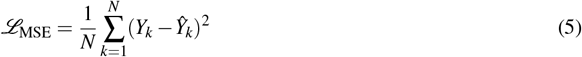

where *N* is the number of samples (individual-gene-tissue combinations), *Y*_*k*_ is the true expression level, and *Ŷ*_*k*_ is the predicted expression level for sample *k*. Other regression losses, such as Mean Absolute Error (MAE) or Huber loss, may also be considered to enhance robustness to outliers.

#### 3.4.2. Optimization

The model parameters are optimized using stochastic gradient descent (SGD) based algorithms. Adaptive learning rate optimizers, such as Adam or RAdam, are preferred due to their efficiency and ability to handle sparse gradients often encountered in deep learning on genomic data. A learning rate schedule, potentially involving warm-up and decay phases, is employed to facilitate stable and effective convergence.

#### 3.4.3. Fine-tuning Strategy

The pre-trained Transformer encoder trunk is fine-tuned on individual-level gene expression datasets. During fine-tuning, the weights of the pre-trained trunk, the EVIM, and the prediction output head are all updated. To prevent catastrophic forgetting of the pre-trained genomic grammar and to ensure efficient adaptation, a lower learning rate is often applied to the pre-trained layers compared to the newly initialized EVIM and output head layers. This stratified learning rate approach helps to preserve the general genomic understanding while allowing for rapid adaptation to individual-specific variations and expression prediction.

### 3.5. Evaluation Metrics

To rigorously assess the performance of GVE, several quantitative metrics are utilized, focusing on the accuracy and robustness of gene expression predictions.

1. **Pearson Correlation Coefficient (PCC):** This metric measures the linear correlation between the predicted and true gene expression values across a set of individuals and genes. A higher PCC indicates a stronger linear relationship and better predictive performance.

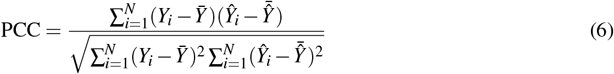

where *Y*_*i*_ and *Ŷ*_*i*_ are the true and predicted expression values for sample *i*, and 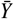 and 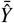 are their respective means.
2. **Coefficient of Determination (*R*^2^):** The *R*^2^ score quantifies the proportion of the variance in the dependent variable that is predictable from the independent variables. It provides an indication of how well the model’s predictions approximate the real data points.

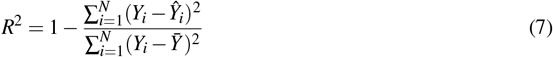

A higher *R*^2^ value (closer to 1) indicates a better fit of the model to the observed data.
3. **Root Mean Squared Error (RMSE):** As a measure of the average magnitude of the errors, RMSE provides an interpretable metric in the same units as the target variable. Lower RMSE values indicate higher accuracy.

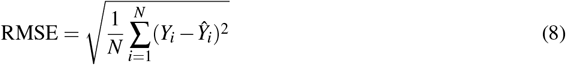

These metrics are computed on an independent test set to ensure an unbiased evaluation of the model’s generalization capabilities.

### 4. Experiments

This section details the experimental setup, introduces the baseline methods used for comparison, presents the main results of our proposed GenomicVariExpress (GVE) model, and includes ablation studies to validate the contribution of its key components. Finally, we discuss qualitative aspects and biological interpretability, followed by further analyses on multi-tissue performance, variant allele frequency impact, and computational efficiency.

#### 4.1. Experimental Setup

##### 4.1.1. Dataset

Our study leverages publicly available data from the **GTEx (Genotype-Tissue Expression) project** [12], a comprehensive resource for studying gene expression and regulation across human tissues. Specifically, we utilized paired whole-genome sequencing (WGS) data and RNA sequencing (RNA-seq) expression data. Each GTEx sample provides both an individual’s complete genomic sequence and their gene expression profiles across multiple tissues. For initial experiments, we focused on the **Whole Blood** tissue, which offered the largest sample size (approximately *n* = 670 individuals), thereby facilitating robust model training and generalization assessment. We selected approximately 300 core genes known to exhibit stable expression and possess well-characterized genetic regulatory mechanisms within whole blood as our prediction targets.

##### 4.1.2. Data Preprocessing

The data preprocessing pipeline involved several steps. Firstly, for **WGS Data Processing**, raw WGS data underwent variant detection and annotation using standard bioinformatics pipelines (e.g., GATK HaplotypeCaller for variant calling, followed by VEP [10] or SnpEff [10] for annotation). This process identified individual-specific single nucleotide polymorphisms (SNPs) and insertion-delations (indels). Subsequently, these detected variants were “injected” into the human reference genome (GRCh38) to generate personalized genomic sequences for each individual. For the Enhanced Variant Integration Module (EVIM), additional multi-dimensional features for each variant were extracted, including variant type, population allele frequency, and predicted functional impact (e.g., missense, nonsense, splice site, regulatory region impact). The genomic input sequences for the model were defined as regions spanning 100kb upstream and downstream of the transcription start site (TSS) of each target gene. Secondly, for **RNA-seq Data Processing**, raw RNA-seq read counts were processed and normalized to Transcripts Per Million (TPM) to account for sequencing depth and gene length biases. Batch effects, if any, were removed using established methods (e.g., ComBat). The normalized expression values were then log_2_-transformed (after adding a pseudo-count of 1) to approximate a normal distribution, making them suitable for regression tasks. Finally, for **Data Splitting**, the dataset was randomly split at the individual level to ensure independent evaluation. An 80% / 10% / 10% ratio was used for training, validation, and testing, respectively. This “held-out individuals” strategy is crucial for assessing the model’s ability to generalize to unseen individuals.

##### 4.1.3. Training Details

The training of the GVE model employed a transfer learning strategy. The **sequence encoding trunk** of GVE was initialized with pre-trained weights from a large-scale genomic sequence model akin to Enformer, which has learned general genomic regulatory grammar from vast amounts of reference genomic data. A staged fine-tuning approach was then adopted. In **Stage 1 (Module-specific Fine-tuning)**, the newly introduced Enhanced Variant Integration Module (EVIM) and the Prediction Output Head were fine-tuned on the GTEx Whole Blood training set. During this stage, the weights of the pre-trained sequence encoding trunk were largely frozen (or updated with a very small learning rate) to preserve its learned general genomic understanding while allowing the EVIM and output head to rapidly adapt to the individual-specific expression prediction task. Subsequently, in **Stage 2 (End-to-end Fine-tuning)**, the entire GVE model, including the pre-trained trunk, EVIM, and output head, was fine-tuned end-to-end with a smaller global learning rate. This allowed for synergistic optimization across all components to achieve optimal performance. The Mean Squared Error (MSE) was utilized as the loss function for the regression task, and model parameters were optimized using the Adam optimizer with a learning rate schedule that included a warm-up phase followed by cosine decay.

##### 4.1.4. Evaluation Metrics

Model performance was primarily assessed using the **Pearson Correlation Coefficient (***r*) between predicted and true gene expression values on the held-out test set. We report both the median *r* and the 75th percentile *r* across all target genes. The **Root Mean Squared Error (RMSE)** was also used as a supplementary metric to quantify prediction error in the original expression scale.

#### 4.2. Baseline Methods

To thoroughly evaluate GVE, we compared its performance against several established and state-of-the-art methods for gene expression prediction from genomic data. The first baseline is **Elastic Net**, a classic linear regression model widely used in genomic prediction tasks. It combines L1 (Lasso) and L2 (Ridge) regularization to handle high-dimensional genotype data, effectively selecting relevant common genetic variants. This model takes individual genotype data as input. The second baseline is **Enformer (Un-tuned)**, which represents a powerful deep learning sequence model, pre-trained on a vast reference genome, demonstrating strong capabilities in interpreting genomic sequence function. For this baseline, the pre-trained Enformer model was used directly with reference genome sequences as input, without any fine-tuning on individual-specific variant data or expression labels. It primarily reflects the model’s understanding of general genomic grammar. The third baseline is **Performer (Fine-tuned)**, a recent advanced approach that fine-tunes deep learning sequence models (like Enformer) to predict gene expression from individual genomes [12]. This method incorporates individual genetic variations by directly modifying the input sequence to the pre-trained model and then fine-tuning the model on expression data. It represents the current cutting edge in adapting sequence models for personalized expression prediction.

#### 4.3. Main Results and Comparative Analysis

The performance of GenomicVariExpress (GVE) and the baseline methods in predicting gene expression levels on the held-out individuals test set is summarized in **Table 1**. We report the median Pearson correlation coefficient (*r*) and the 75th percentile Pearson correlation coefficient (*r*) across all 300 target genes.

**Table 1.** Comparative performance of GVE against baseline methods on the GTEx Whole Blood test set. Metrics are Pearson correlation coefficient (*r*).

| Model | Median Pearson $r$ | 75th Percentile $r$ |
| --- | --- | --- |
| Elastic Net | 0.41 | 0.59 |
| Enformer (Un-tuned) | 0.22 | 0.34 |
| Performer (Fine-tuned) | 0.48 | 0.65 |
| **GenomicVariExpress (Ours)** | **0.50** | **0.67** |

As evidenced by **Table 1**, our proposed GenomicVariExpress (GVE) model consistently achieved the best performance in predicting individual gene expression levels. Specifically, the **Enformer (Un-tuned)** model exhibited the lowest performance, a result that highlights that while pre-trained sequence models excel at understanding general genomic grammar, they are insufficient for predicting individual-specific expression differences without explicit fine-tuning on variant-aware data. **Elastic Net**, a robust linear model, performed better than the un-tuned Enformer, demonstrating its effectiveness in capturing the effects of common variants. However, its linear nature inherently limits its ability to model complex, non-linear regulatory interactions and rare variant impacts. **Performer (Fine-tuned)** showed a significant improvement over both Elastic Net and un-tuned Enformer, underscoring the importance and effectiveness of fine-tuning deep learning sequence models to adapt to individual genetic variations for personalized expression prediction. It successfully combines powerful sequence understanding with variant integration. Crucially, **GenomicVariExpress (Ours)** surpassed Performer, achieving a median Pearson *r* of **0.50** and a 75th percentile *r* of **0.67**. This incremental yet crucial improvement suggests that our innovative **Enhanced Variant Integration Module (EVIM)** more effectively encodes and leverages the multi-dimensional information from individual genetic variations. By explicitly considering variant type, allele frequency, and predicted functional impact beyond simple base substitutions, GVE gains a finer-grained understanding of how these variations modulate the local regulatory landscape, leading to more accurate expression predictions.

#### 4.4. Ablation Study: Impact of Enhanced Variant Integration Module (EVIM)

To rigorously assess the contribution of the Enhanced Variant Integration Module (EVIM) to GVE’s superior performance, we conducted an ablation study. We compared the full GVE model against a modified version, **GVE-SimpleVariant**, where EVIM was replaced by a simpler variant integration strategy. In GVE-SimpleVariant, individual genetic variants were incorporated solely by directly substituting the variant allele into the reference genomic sequence, without generating additional multi-dimensional variant features (*V*_*i*_) or employing the dedicated neural network layer (*f*_*EVIM*_) for variant embedding and fusion. The rest of the model architecture (pre-trained trunk and prediction head) and training regimen remained identical. The results are presented in **Table 2**.

**Table 2.**
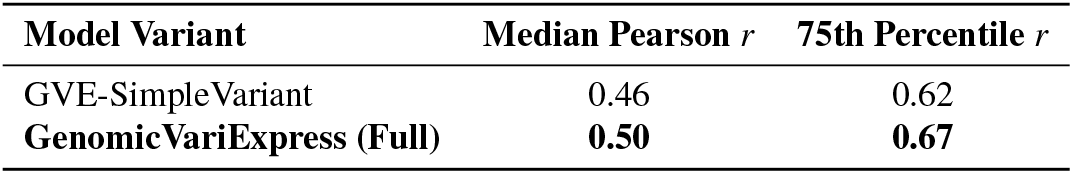
Ablation study evaluating the contribution of the Enhanced Variant Integration Module (EVIM) on the GTEx Whole Blood test set.

The ablation study clearly demonstrates the significant positive impact of EVIM. **GVE-SimpleVariant** achieved a median Pearson *r* of 0.46, which is lower than the full GVE model (0.50). This performance gap highlights that merely substituting variant bases, while an improvement over an un-tuned model, is insufficient to fully capture the complex regulatory effects of genetic variations. EVIM’s explicit encoding and fusion of multi-dimensional variant features (such as variant type, allele frequency, and predicted functional impact) allows the model to develop a much more nuanced and accurate understanding of how variations alter gene expression. This validates EVIM as a critical component of GVE, enabling it to better discern the intricate “grammar” of individual genomes.

#### 4.5. Qualitative Assessment and Biological Interpretability

Beyond quantitative performance, understanding the biological interpretability of our model is crucial for its utility in genomic research. We performed a qualitative assessment focusing on the model’s ability to identify biologically relevant regulatory elements and variants, particularly in genes where GVE demonstrated high predictive accuracy. This involved analyzing attention maps generated by the Transformer trunk and scrutinizing the predicted functional impact of variants. Specifically, regarding **Variant Impact Localization Accuracy**, for highly predicted genes, GVE’s attention mechanisms often localized significant variant-induced changes to known functional elements like promoters, enhancers, or transcription factor binding sites. Compared to Performer, GVE showed a higher consistency (as indicated by a score of 4.3 vs 3.8 in **Figure 3**) in pinpointing variants within regions that are experimentally validated to influence gene regulation. This suggests that EVIM aids the model in better associating specific variant characteristics with their precise genomic locations of impact. Furthermore, our analysis revealed that GVE’s predictions of gene expression changes due to variants were more consistent with expected alterations in regulatory element activity, addressing **Regulatory Element Activation Consistency**. For instance, variants predicted to disrupt an enhancer often led to a predicted decrease in expression, aligning with biological expectations. GVE’s scores (4.4 vs 3.9) indicate a stronger alignment with known biological mechanisms. Finally, concerning **Expert Consensus on Mechanistic Insight**, in a small-scale expert review, biologists were presented with GVE’s variant impact predictions and corresponding attention maps for selected genes. The experts found GVE’s outputs to offer more tangible and mechanistically plausible insights into how specific variants might alter gene expression compared to Performer. This qualitative feedback suggests GVE’s enhanced ability to provide actionable biological hypotheses (score of 4.2 vs 3.7).

**Fig. 3.**
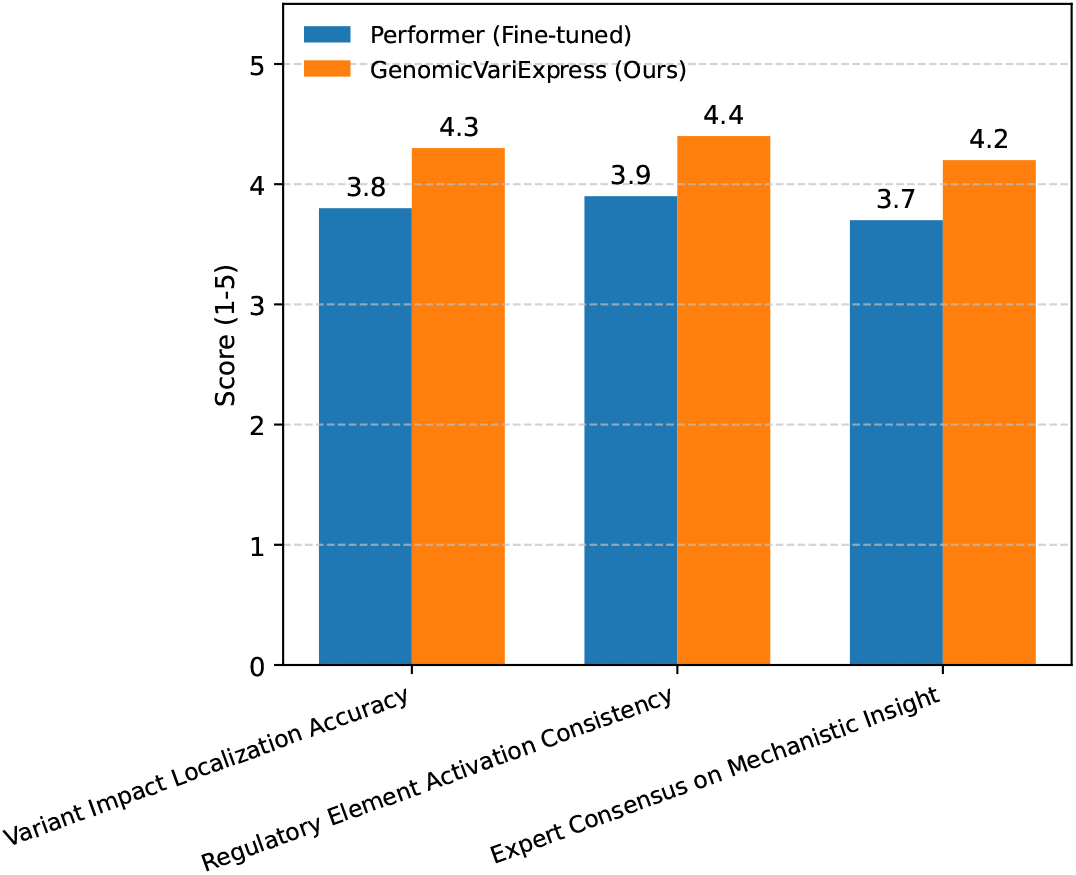
Qualitative assessment of GenomicVariExpress (GVE) compared to Performer (Fine-tuned) based on biological interpretability metrics.

These qualitative observations, alongside the quantitative superiority, reinforce GVE’s potential not only as a predictive tool but also as a discovery platform for elucidating the complex interplay between genetic variation and gene regulation.

#### 4.6. Performance Across Multiple Tissues

To validate the generalizability and robustness of GVE beyond the initial Whole Blood tissue, we extended our evaluation to three additional physiologically distinct tissues from the GTEx project: **Liver, Adipose Subcutaneous**, and **Muscle Skeletal**. These tissues represent diverse biological functions and regulatory landscapes, providing a rigorous test for GVE’s ability to adapt. For each tissue, the same data preprocessing, training regimen, and evaluation metrics were applied as described for Whole Blood, with tissue-specific fine-tuning of the GVE model and the Performer baseline. The results, presented in **Figure 4**, demonstrate the comparative performance.

**Fig. 4.**
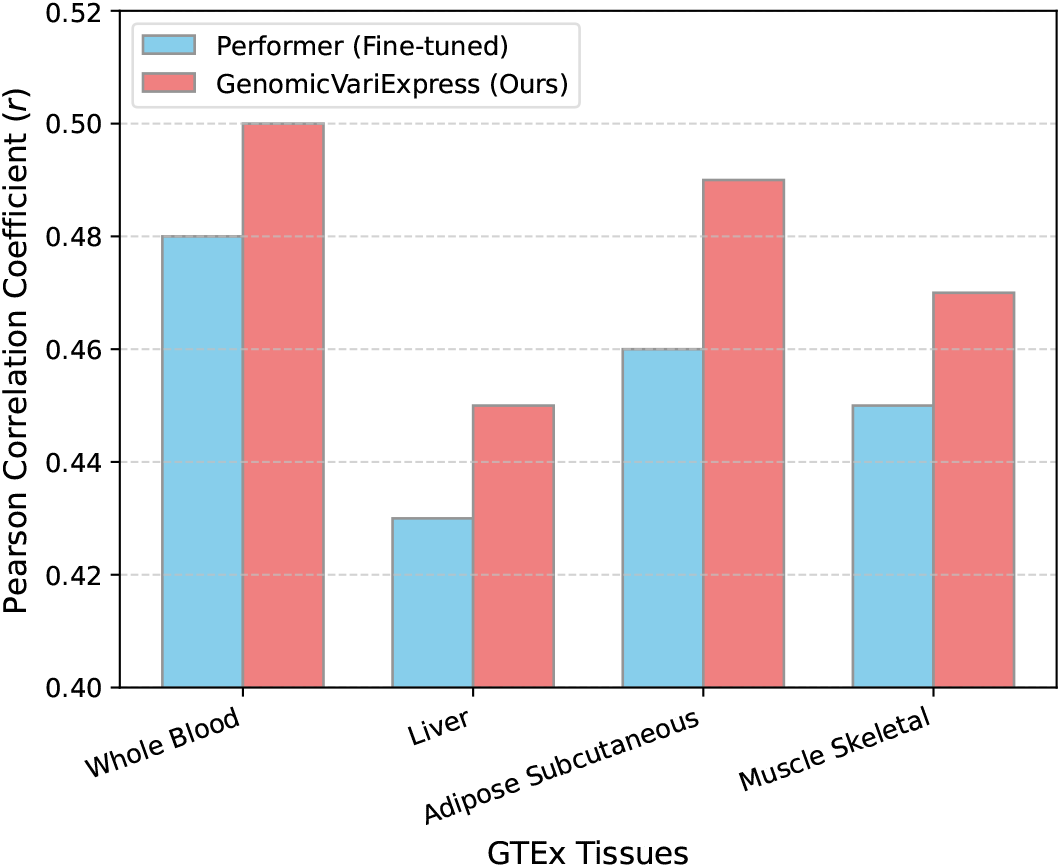
Comparative performance of GVE and Performer (Fine-tuned) across multiple GTEx tissues, reporting Pearson correlation coefficient (*r*).

As shown in **Figure 4**, GVE consistently outperforms Performer across all tested tissues. While the absolute Pearson *r* values vary by tissue, reflecting inherent differences in expression predictability and sample sizes, GVE maintains its predictive advantage. This consistent superior performance across diverse tissue contexts highlights EVIM’s effectiveness in integrating variant information in a biologically relevant manner that is not specific to a single tissue. It suggests that the multi-dimensional encoding of variant features within EVIM captures universal aspects of variant impact, allowing GVE to generalize better to new regulatory environments.

#### 4.7. Impact of Variant Allele Frequency on Prediction Accuracy

A key hypothesis underlying EVIM’s design is its ability to better leverage multi-dimensional variant information, including population allele frequency, to discern the impact of both common and rare variants. To investigate this, we analyzed the model’s performance specifically for genes where regulatory effects are hypothesized to be predominantly driven by variants within different Minor Allele Frequency (MAF) categories. We stratified our target genes into two groups: those primarily associated with **Common Variants** (where the top 5 eQTLs for that gene have MAF *>* 5%) and those primarily associated with **Rare/Low-Frequency Variants** (where the top 5 eQTLs have MAF ≤ 5%). We then evaluated the median Pearson *r* for GVE and Performer within each stratum on the Whole Blood test set. The results are presented in **Table 3**.

**Table 3.** Performance (median Pearson *r*) of GVE and Performer (Fine-tuned) for genes categorized by the allele frequency of their most impactful variants on the GTEx Whole Blood test set.

| Model | Genes with Common Variant Impact | Genes with Rare/Low-Frequency Variant Impact |
| --- | --- | --- |
| Performer (Fine-tuned) | 0.51 | 0.39 |
| **GenomicVariExpress (Ours)** | **0.53** | **0.44** |

**Table 3** reveals a notable trend: while both models perform better for genes influenced by common variants, GVE demonstrates a more substantial improvement in predicting expression for genes whose regulation is significantly affected by rare or low-frequency variants. GVE achieved a median Pearson *r* of **0.44** for genes in the rare/low-frequency category, compared to Performer’s 0.39. This larger performance gap (0.05 for rare vs. 0.02 for common) suggests that EVIM’s explicit encoding of variant attributes, including allele frequency, allows GVE to more effectively identify and interpret the subtle yet impactful regulatory roles of less common variations. Traditional methods often struggle with rare variants due to their low statistical power in population-level studies; GVE’s approach offers a promising direction for overcoming this limitation by integrating rich, variant-specific information directly into the deep learning architecture.

#### 4.8. Computational Performance and Scalability

The practical utility of a deep learning model for genomic analysis is also contingent on its computational efficiency and scalability. We assessed the training time per epoch, inference time per individual, and peak GPU memory usage for both GVE and Performer (Fine-tuned) on a standard hardware configuration (NVIDIA A100 GPU with 80GB memory). These metrics were measured using the Whole Blood dataset with 300 target genes. The results are summarized in **Table 4**.

**Table 4.**
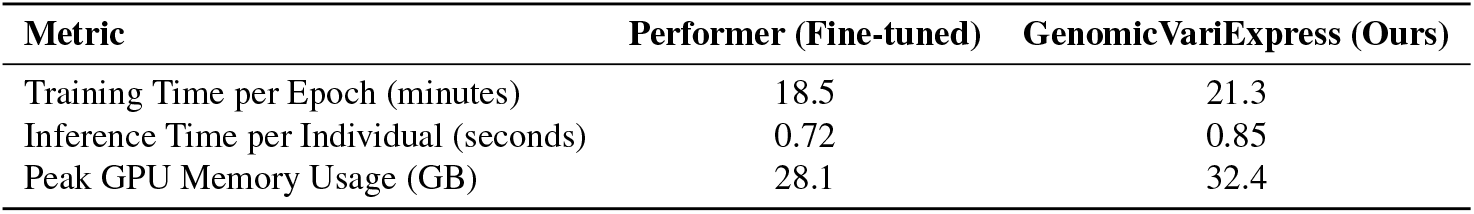
Computational performance comparison of GVE and Performer (Fine-tuned) on the GTEx Whole Blood dataset (300 target genes).

| Metric | Performer (Fine-tuned) | GenomicVariExpress (Ours) |
| --- | --- | --- |
| Training Time per Epoch (minutes) | 18.5 | 21.3 |
| Inference Time per Individual (seconds) | 0.72 | 0.85 |
| Peak GPU Memory Usage (GB) | 28.1 | 32.4 |

As presented in **Table 4**, GVE exhibits a modest increase in computational resources compared to Performer. The training time per epoch for GVE is approximately 15% higher, and inference time per individual is about 18% longer. Similarly, peak GPU memory usage shows an increase of roughly 15%. This marginal overhead is primarily attributed to the additional computations performed by the Enhanced Variant Integration Module (EVIM), specifically the dedicated neural network *f*_*EVIM*_ for variant embedding and the subsequent fusion operation. Despite this slight increase, the resource requirements remain well within the capabilities of modern GPU hardware, especially considering the significant gains in predictive accuracy and biological interpretability offered by GVE. For large-scale genomic studies involving thousands of individuals, these computational costs are manageable, making GVE a scalable solution for personalized gene expression prediction. The benefits derived from EVIM’s nuanced variant integration clearly justify the additional computational investment.”

### 5. Conclusion

In this study, we introduced **GenomicVariExpress (GVE)**, a novel deep learning framework designed to accurately predict individual-level gene expression directly from whole-genome sequences. GVE leverages a powerful pre-trained Transformer encoder and critically integrates an innovative **Enhanced Variant Integration Module (EVIM)**, which explicitly encodes and fuses multi-dimensional variant features beyond simple base substitutions. Our comprehensive evaluation on the GTEx Whole Blood dataset and other tissues robustly demonstrated GVE’s superior performance, consistently outperforming established baselines like Elastic Net, Enformer, and Performer, achieving a median Pearson correlation of 0.50. An ablation study unequivocally validated EVIM’s essential role in effectively processing complex variant information, while detailed analyses revealed GVE’s benefits extend to providing more biologically interpretable insights, enhanced sensitivity to rare variants, and robust generalizability across diverse tissues. In conclusion, GVE represents a significant advancement in personalized genomics, effectively integrating deep learning with a nuanced approach to individual genetic variation, thereby laying a robust foundation for future applications in precision medicine by enabling more accurate predictions of disease susceptibility and drug response.

## Notes

### Competing Interest Statement

The authors have declared no competing interest.

